# 3D architecture and a bi-cellular mechanism of touch detection in mechanosensory corpuscle

**DOI:** 10.1101/2023.04.05.535701

**Authors:** Yury A. Nikolaev, Luke H. Ziolkowski, Song Pang, Wei-Ping Li, Viktor V. Feketa, C. Shan Xu, Elena O. Gracheva, Sviatoslav N. Bagriantsev

## Abstract

Mechanosensory corpuscles detect transient touch and vibratory signals in the skin of vertebrates, enabling navigation, foraging, and precise manipulation of objects^1^. The corpuscle core comprises a terminal neurite of a mechanoreceptor afferent, the only known touch-sensing element within corpuscles, surrounded by terminal Schwann cells called lamellar cells (LCs)^2–4^. However, the precise corpuscular ultrastructure, and the role of LCs in touch detection are unknown. Here we used enhanced focused ion beam scanning electron microscopy and electron tomography to reveal the three-dimensional architecture of avian Meissner (Grandry) corpuscle^5^. We show that corpuscles contain a stack of LCs innervated by two afferents, which form large-area contacts with LCs. LCs form tether-like connections with the afferent membrane and contain dense core vesicles which release their content onto the afferent. Furthermore, by performing simultaneous electrophysiological recordings from both cell types, we show that mechanosensitive LCs use calcium influx to trigger action potential firing in the afferent and thus serve as physiological touch sensors in the skin. Our findings suggest a bi-cellular mechanism of touch detection, which comprises the afferent and LCs, likely enables corpuscles to encode the nuances of tactile stimuli.

## Main Text

The sense of touch is essential for handling objects and tools, foraging, navigating an environment, and forming social bonds^6^. In vertebrates, the various properties of touch are detected in the skin by mechanosensory corpuscles and hair follicle-associated lanceolate complexes^1, 7^. Although morphologically and functionally diverse, these end organs invariably contain terminal Schwann cells (TSC) that form close interactions with mechanoreceptor afferent terminals. The afferents, which express mechanically gated ion channels such as Piezo2^8, 9^, are considered to be the only touch-sensing elements within these end organs, but recent work has revealed that dissociated mouse Schwann cells display mechanosensitivity in culture^10, 11^. Moreover, mammalian Meissner corpuscles and their avian analogs (historically called corpuscles of Grandry and referred to herein as avian Meissner corpuscles) contain TSCs known as lamellar cells (LC)^2, 5^, which fire mechanically activated action potentials in intact avian Meissner corpuscles *in situ*^3^. These studies suggest that the sensory afferent may not be the sole sensory element, and that LCs could be additional physiological sensors of touch. However, the detailed architecture of LC-afferent complexes is unknown. Thus, the role of LCs remains speculative in the absence of structural and functional insight into the relationship between lamellar cells and sensory afferents^12^.

### Architecture of avian Meissner corpuscles

To determine the detailed structure of a Meissner corpuscle, we used focused ion beam scanning electron microscopy (FIB-SEM) to image bill skin from a tactile foraging Mallard duck (*Anas platyrhynchos domesticus*). We scanned a volume of 47,250 μm^3^, comprising 4,753 consecutive images at a voxel size of 8 x 8 x 8 nm, and used machine learning to reconstruct the 3D architecture of an entire LC-afferent complex and its associated structures. The boundary box for reconstruction had dimensions of 35 x 45 x 30 μm, of which, the corpuscle occupied a volume of 8,167 μm^3^ (Fig. 1A, Supplementary Table I).

**Fig. 1.**
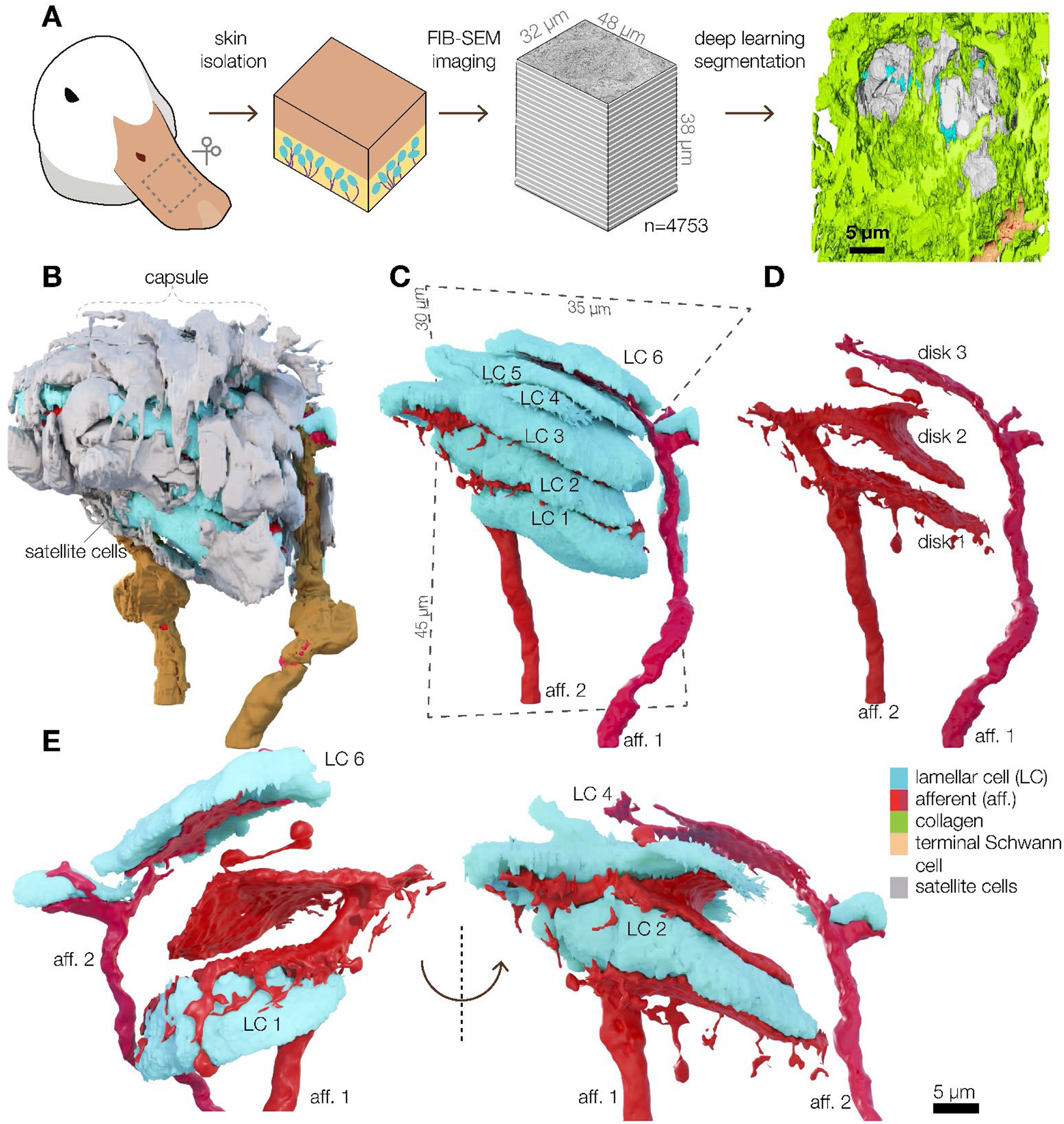
Meissner corpuscles comprise a stack of LCs interdigitated with terminal afferent disks. **(A)** FIB-SEM workflow for automated segmentation and machine learning-based 3D reconstruction of a Meissner corpuscle in duck bill skin. **(B**—**D)** 3D architecture of a Meissner corpuscle (B), corpuscle without outer capsule (C), isolated afferents (E). **(E)** 3D architecture of a section of afferent 1 and afferent 2 and associated LCs.

The outer layer of the corpuscle, formed by satellite cells and collagen fibers, encapsulates a sensory core comprising a stack of six LCs (LC1-6) innervated by two afferents (Fig. 1B-D, Movie 1). Both afferents are covered with myelinating Schwann cells outside the corpuscles (Fig. 1B, C). Afferent 1 interdigitates with the LCs, forming disk-like endings that cover up to 42% of the apposing surfaces of LC1-6 (Fig. 1D, E, Extended Data Fig. 1). The discoid endings form protrusions that extend around LC1 (Fig. 1E). Afferent 2 forms a single, smaller, ovoid ending between LC5 and LC6 and covering 16% of the lower surface of LC6 (Fig. 1C-E, Extended Data Fig. 1). One of the satellite cells, whose cell body was outside of the afferent-LC core, formed fine projections interleaved between afferent 2 and LC5 (Extended Data Fig. 2). We reconstructed the LC-afferent core of another corpuscle from the same skin volume, which contained a stack of 3 LCs. This corpuscle is innervated by a single afferent which, like afferent 2, formed a single disc in the LCs stack (Extended Data Fig. 3). Thus, avian Meissner corpuscles can be innervated by one or two afferents. The difference in the innervation pattern among the afferents indicates they could be molecularly and physiologically distinct^2^.

### Structural coupling between lamellar and afferent membranes

Between the disk-shaped afferents, the six LCs form flattened structures, 6-9.6 μm thick and 23.9-33.1 μm along the longest dimension. Small villi protrude from the edges of each LC and form contacts with the surrounding satellite cells and collagen (Fig. 2A, B). The cytosol of each LC contains 19,981-37,306 dense core vesicles (DCVs), ∼150 nm in diameter, which occupy 2-3% of the volume of each LC (Fig. 2B-E, Extended Data Fig. 4). Approximately 1% of all DCVs were within 30 nm from the membrane facing the afferent, suggesting they could represent a ready-releasable pool (Fig. 2F). We used transmission electron microscopy and electron tomography to reconstruct and segment a 3.16 x 2.16 x 0.25 μm box containing an LC-afferent disk contact (Video 2, Video 3). A high-resolution close-up reconstruction of a fragment of the LC-afferent contact area revealed that DCVs fuse with regions of LC plasma membrane that appose the neuronal disk, suggesting that DCVs release their contents into the intermembrane space (Fig. 2G, H, Video 4). Densely coated caveolae were also found adjacent to DCVs (Fig. 2G, H, Video 4). RNA sequencing of corpuscles extracted from bill skin revealed robust expression of components of DCV biogenesis and release, including SNARE proteins, as well as mechanically gated and voltage-gated ion channels (Extended Data Fig. 5). This supports our observation of DCVs and earlier data showing that LCs are mechanosensitive and excitable^3^.

**Fig. 2.**
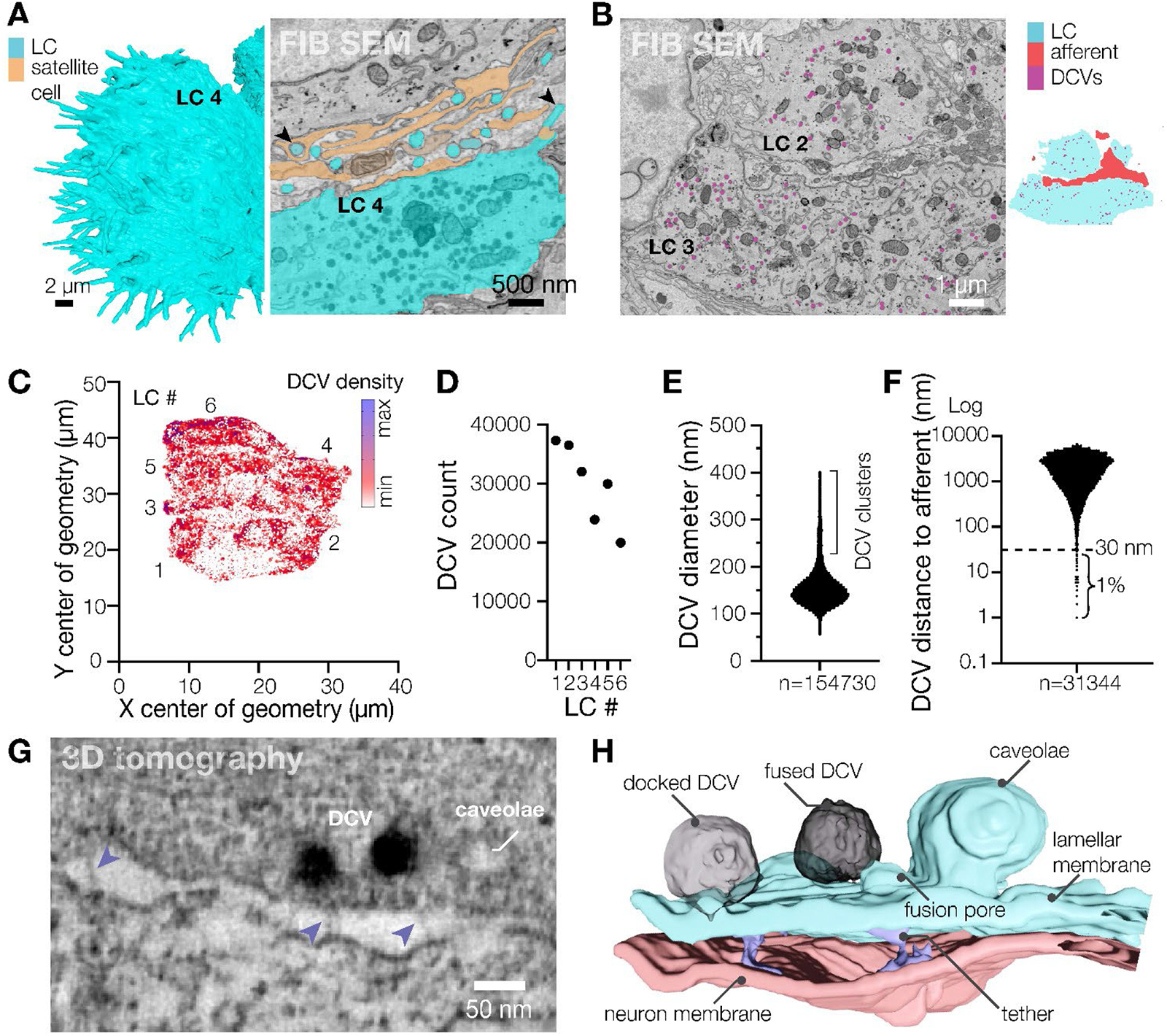
Lamellar cells interact with the afferent via DCVs and tethers. **(A)** Close-up 3D reconstruction of villi protruding from the edge of LC4 and a pseudo-colored scanning electron microscope image of villi (black arrowheads) protruding from the edge of LC4 and contacting the satellite cell and afferent. **(B)** A scanning electron microscope image depicting DCVs inside LCs (left) and a corresponding map of the cell types shown in the image (right). **(C)** A density map of DCVs in LCs. **(D-F)** Quantification of the DCV number (D), diameter (E) and distance to the afferent (F). **(G)** Transmission electron microscopy image of the LC–afferent contact area. Blue arrowheads point to tethers connecting LC and afferent plasma membranes **(H)** 3D reconstruction of a fragment of the LC-afferent contact area.

Although the sensory afferent in the reconstruction contained smaller, clear vesicles, we did not detect synapse-like structures between the LC and afferent plasma membranes, but found membrane densities resembling adherens junctions (Video 2, Video 3). Additionally, we observed 26-32 nm-long tethers connecting the membranes throughout LC-afferent contact area, including at sites of DCV fusion (Fig. 2G, H, Video 4). Together with earlier data^3^, these findings reveal that avian Meissner LCs are mechanosensitive secretory cells that form large contact areas with sensory afferents and tether-like connections with afferent membranes, suggesting possible functional interaction between LCs and afferent terminals.

### Mechanosensitive lamellar cells activate the afferent

Because patch-clamp recordings directly from Meissner LCs in duck bill skin previously revealed that LCs are mechanosensitive^3^, we sought to test the idea that LCs can induce firing in sensory afferents and thereby act as *bona fide* touch sensors. As a first step, we used a method for electrophysiological recordings of afferent nerve activity from a single corpuscle in duck bill skin during mechanical stimulation of the same corpuscle (Fig. 3A)^13^. Mechanical stimulation of the corpuscle with a blunt glass probe mounted on a piezoelectric actuator evoked action potential (AP) firing in the afferent (Fig. 3B). APs occurred in a 1:1 correspondence with indentations during a 20 Hz vibratory stimulation (Fig. 3D), consistent with the role of avian Meissner corpuscles as detectors of transient touch and vibration^5, 14, 15^. We detected two types of firing pattern in the afferent. In most cases, APs were triggered in the rapidly adapting fashion, i.e. only during the dynamic phases of the stimulus (Fig. 3E). However, two out of more than 50 recorded afferents showed slowly adapting firing (Extended Data Fig. 6). This supports with our structural data showing two morphologically distinct afferents in the corpuscle and agrees with earlier findings in mice^2^. Application of the voltage-gated sodium channel blocker tetrodotoxin (TTX) suppressed AP generation, revealing low-amplitude receptor potentials (Fig. 3B-D), consistent with the presence of mechanically gated ion channels in the afferent ending^13^.

**Fig. 3.**
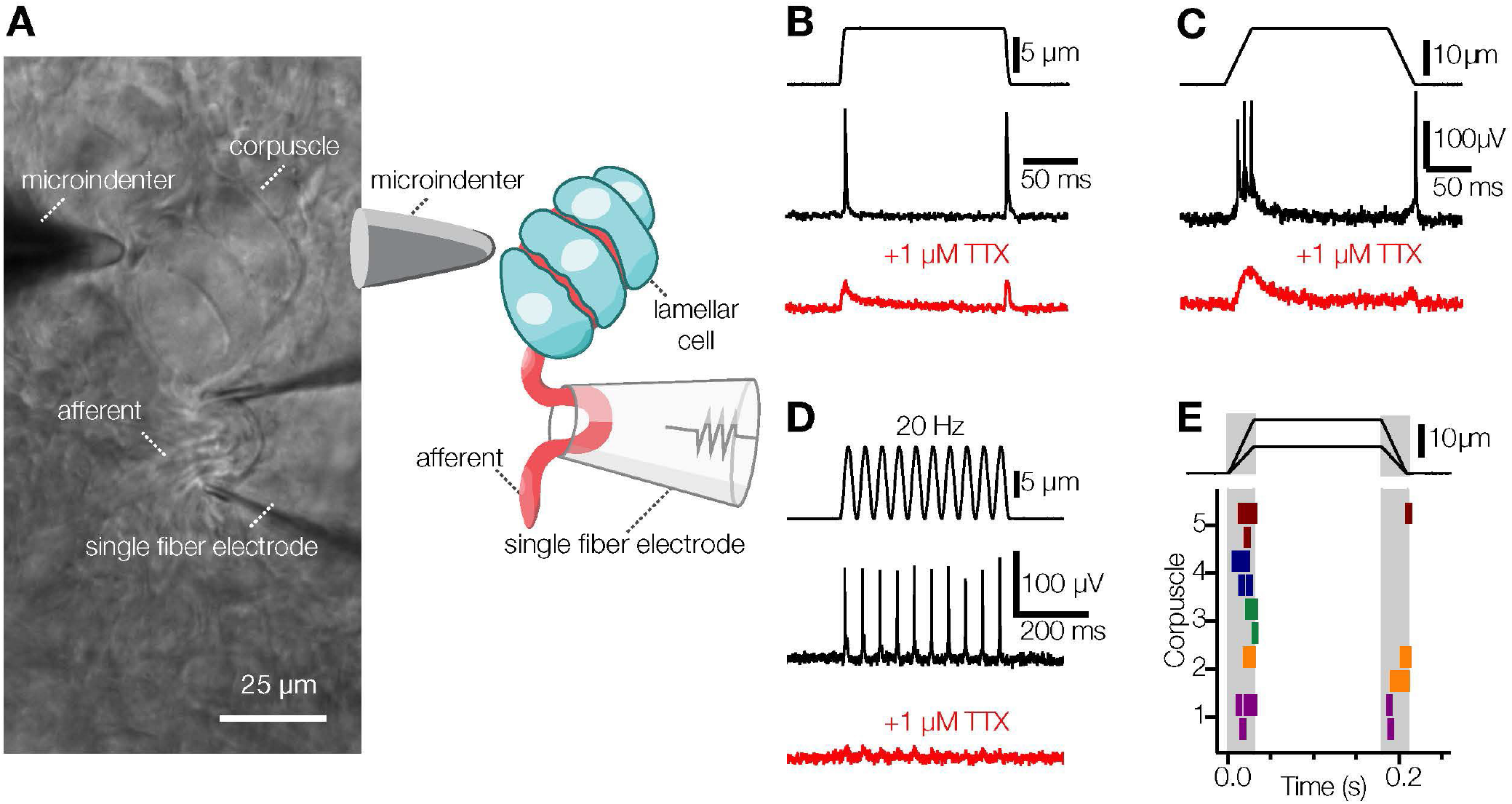
Avian Meissner corpuscles detect transient touch. **(A, B)** Experimental setup visualized as a bright-field image. (A) and schematic representation (B). **(C)** Mechanical step stimulus applied with a glass probe (top), representative rapidly adapting single-fiber response comprising APs (middle), and representative single-fiber response in 1 μM TTX comprising receptor potentials (bottom). **(D)** Mechanical step stimulus with long ramp phases (top), representative rapidly adapting single-fiber response comprising APs (middle), and representative single-fiber response in 1 μM TTX comprising receptor potentials (bottom). **(E)** Vibratory mechanical stimulus (top), representative single-fiber response comprising APs (middle), and representative single-fiber response in 1 μM TTX comprising receptor potentials (bottom). **(F)** Raster plot of rapidly adapting afferent firing for five different corpuscles (bottom) in response to mechanical stimuli of two different indentation depths. Each vertical dash represents an individual AP.

Mechanical stimulation of the corpuscle inevitably acts on both the afferent and LCs, therefore, to test whether activation of a single LC can trigger afferent firing, we selectively activated a single LC by whole-cell patch-clamp while simultaneously recording afferent activity from the same corpuscle with a second electrode (Fig. 4A, B). To our knowledge, such an experiment has not been performed for any tactile corpuscle in any species. When LCs were stimulated by depolarizing current injection in the current-clamp mode, they displayed robust AP firing, as expected^3^. Interestingly, concurrent AP firing was also detected in the recorded afferents (Fig. 4C, E). The waveforms in the afferent triggered by LC activation were indistinguishable from the waveforms triggered by mechanical stimulation of the corpuscle (Extended Data Fig. 7A, B). Next, we tested if APs can be triggered in the afferent when LCs are activated by voltage-clamped depolarization to 0 mV, which leads to calcium influx and potassium efflux from LCs^3^. This type of LC activation also led to robust AP firing (Fig. 4D, E, G, Extended Data Fig. 7C, D). The onset of afferent firing was variable, commencing 87-288 ms after the start of current injections and 20-9,120 ms after the start of depolarizations, with an apparent trend towards reduced number of APs towards the end of the stimulus (Fig. 4F). Thus, activation of a single LC is sufficient to drive AP firing in the afferent, demonstrating that LCs can transmit touch information to the afferent.

**Fig. 4.**
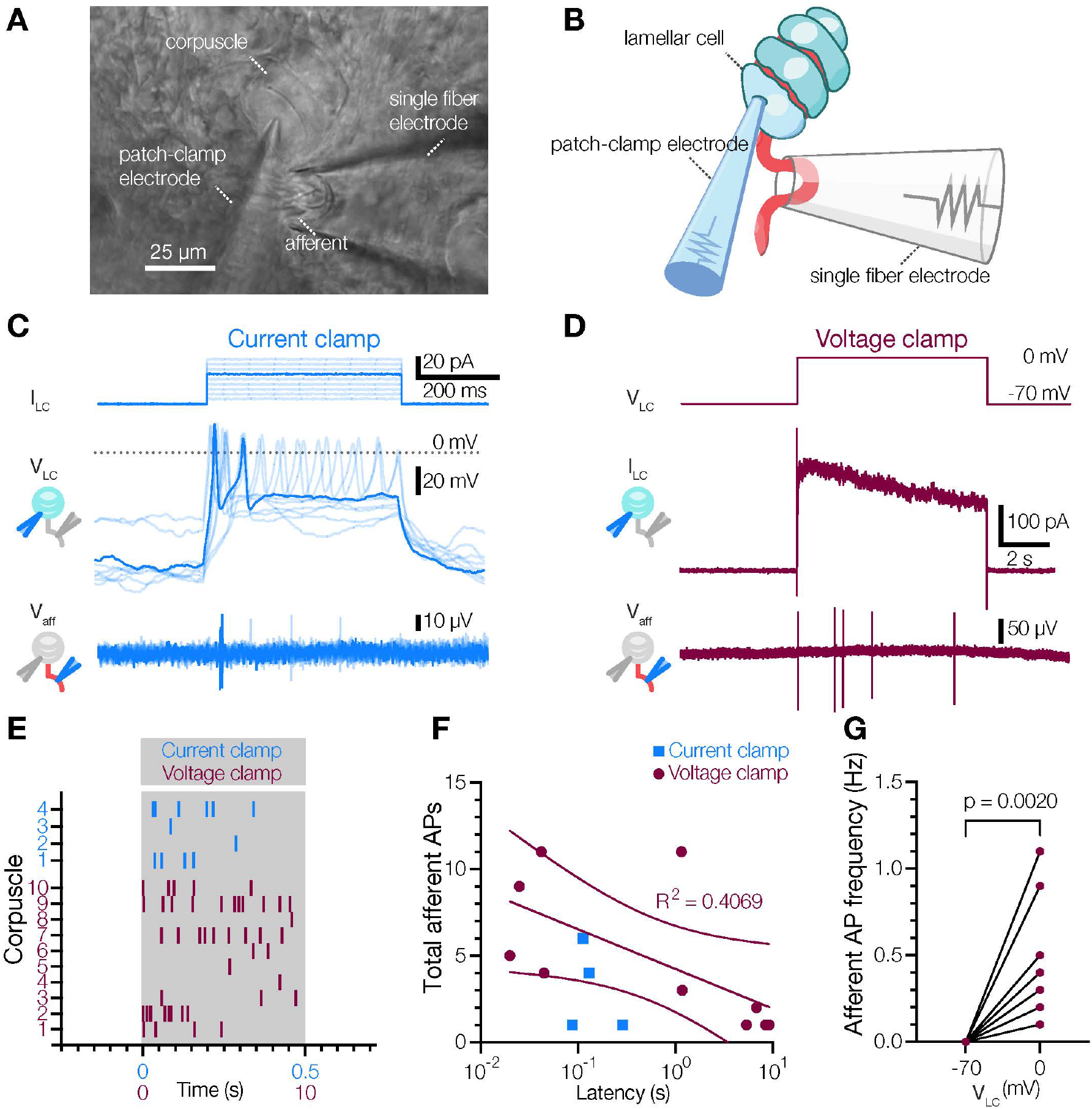
Activation of a single LC is sufficient to drive afferent firing. **(A)** Bright-field image of the experimental setup. **(B)** Schematic representation of the experimental setup. **(C)** Current injection applied to the LC (I_LC_, top), voltage response and APs in the LC (V_LC_, middle), and extracellular voltage and APs in the afferent (V_aff_, bottom). **(D)** Voltage step stimulus applied to the LC (V_LC_, top), current response with potassium-based internal in the LC (I_LC_, middle), and extracellular voltage and APs in the afferent (V_aff_, bottom). **(E)** Raster plot of afferent AP firing for individual corpuscles during LC activation by either depolarizing current injections in current clamp (blue), or voltage steps to 0 mV in voltage clamp (maroon). Each vertical dash represents an individual AP. **(F)** Total number of afferent APs versus latency between onset of stimulus and first AP. Solid line is a fit to the liner equation, dashed lines are 95% confidence intervals of the linear fit **(G)** Afferent AP frequency when LCs held at -70 mV and 0 mV. Wilcoxon matched-pairs test.

Next, we sought to probe the mechanism of LC-afferent communication. Because mechanical stimulation of LCs leads to AP firing via voltage-gated calcium channels^3^, we hypothesized that such mechanism could depend on calcium entry into LC. Because depolarization of LCs activates both voltage-activated calcium and voltage-activated potassium channels, we included cesium chloride in the intracellular solution to quench potassium efflux. Under these conditions, activation of LCs successfully induced AP firing in the afferent (Fig. 5A). However, the removal of calcium from the extracellular medium and blockade of voltage-gated calcium channels suppressed LC-induced firing in the afferent, and the effect was reversed by calcium re-introduction (Fig. 5A, B). These data demonstrate that calcium influx during LC activation is an essential prerequisite for LC-afferent communication.

**Fig. 5.**
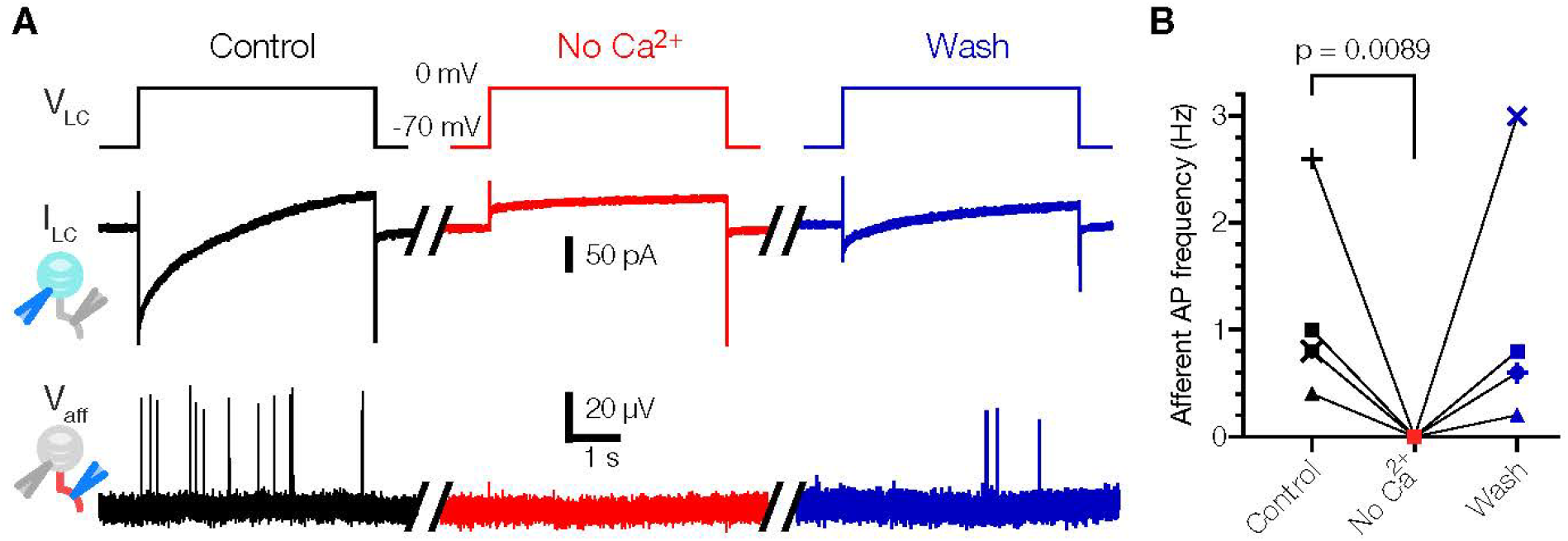
LC activation triggers afferent firing via a calcium-dependent mechanism. **(A)** Exemplar traces recorded in a Meissner afferent in response to LC activation by voltage-clamp with cesium-based internal (Control), following the removal of extracellular Ca^2+^ and addition of 300 μM Cd^2+^ to block voltage-activated calcium channels (No Ca^2+^), and upon reintroduction of calcium and removal of Cd^2+^ (Wash). V_LC_, voltage step stimulus applied to the LC; I_LC_, current response in the LC; V_aff_, extracellular voltage and APs in the afferent. **(B)** Quantification of the effect of calcium removal on LC-induced afferent firing. Lines connect data from the same afferent (n = 5). The difference between Control and Wash was not significant (p = 0.6856). Friedman test with Dunn’s test for multiple comparisons.

## Discussion

Together with earlier observations that avian Meissner LCs are mechanosensitive and excitable^3^, these results establish LCs as physiological sensors of touch. Activation of LCs did not result in immediate excitability changes in the afferent, but instead induced irregular firing with variable delays, arguing against direct electrical coupling. This is consistent with the earlier finding that LCs lack functional gap junctions with surrounding cells^3^. The presence of exocytotic machinery in LCs instead suggests that communication with afferent fibers may involve chemical transmission^16, 17^. In support of this idea, the removal of extracellular calcium abolished LC-induced firing in the afferent, suggesting a possible involvement of calcium-dependent vesicular communication. Other mechanisms, including ephaptic crosstalk, may also be employed. Tethers connecting the membranes of LCs and afferents could provide another alternative, mediating mechanical coupling between cellular elements and/or gating of mechanically activated ion channels^4, 18, 19^. These tethers may also have important biomechanical consequences for the transduction of touch within the corpuscle.

Our findings provide support for a new model of touch detection in Meissner corpuscles by the afferent and LCs. The physiological consequences of a bi-cellular mechanism to detect touch are intriguing. Although mechanoreceptive afferents faithfully fired APs during dynamic stimulation of corpuscles, LCs induced afferent firing with latencies ranging from 20 ms to up to ∼9 s. Thus, LCs are likely not responsible for the fast primary mechanosensory response observed in the afferent^13^. Rather, it is expected that sensory afferents directly mediate this response via Piezo2^8, 20, 21^, and that LCs perform a yet unidentified but complementary role in touch detection. Such a bi-cellular mechanism, which includes the afferents and LCs, would enable multifaceted mechanosensation, potentially facilitating more versatile and precise detection of touch.

Avian Meissner corpuscles are structurally and functionally similar to their mammalian counterparts. Meissner corpuscles endow humans with the remarkable ability to manipulate objects and tools, and enable tactile specialist birds to carry out a multitude of foraging behaviors^5, 7, 22–25^. Whether mammalian LCs are mechanosensitive remains to be determined, but appears likely given that dissociated murine Schwann cells are mechanosensitive in culture^10, 11^. Moreover, there is functional homogeneity between avian and mammalian Meissner corpuscles, suggesting that the physiological role of LCs is conserved between species. While this manuscript was finalized, a preprint study reported the 3D architecture of mouse Meissner corpuscle, which, together with earlier literature, permits a detailed comparison with the avian structure^26^. Both corpuscles detect the same stimuli – transient touch and low-frequency vibration, and contain a stack of LCs innervated by one or two afferents which are mostly, though not exclusively, rapidly adapting^1, 2, 4, 5, 13–15, 23, 26^. At the same time, some differences exist between the two. Unlike mammalian LCs, avian LCs do not form extensive lamellae. Additionally, DCVs are abundantly present in avian LCs, but have not been reported in mammalian Meissner LCs^2, 26^. Interestingly, however, LCs from both avian and mammalian Meissner corpuscles are of mesenchymal origin, similar to the inner core cells in Pacinian corpuscles but dissimilar to epidermal Merkel cells^27–30^. This suggests that other vertebrate end organs in which TSCs are in close association with afferent fibers, including Pacinian corpuscles and hair follicles, may utilize TSCs to detect tactile stimuli^7^.

## Acknowledgements

We thank members of the Bagriantsev and Gracheva laboratories for comments and critique throughout the study; Morven Graham, Xinran Liu and Yale School of Medicine Electron Microscopy Core for transmission electron microscopic imaging for electron tomography; David Ginty for sharing the skin sample fixation protocol; Sue Ann Mentone for help with skin tissue fixation; Güneş Parlakgül, Andrew Bergen and Mariia Burdyniuk for advice on FIB-SEM data processing; Benjamin Bae and Samuel Bae for help with electron microscopy image processing for 3D reconstruction.

## Author contributions

Conceptualization: YAN, LHZ, EOG, SNB. Data collection: YAN, LHZ, SP, DM-A, CSX. Funding acquisition, project administration and supervision: CSX, EOG and SNB. Writing: YAN, LHZ, EOG, SNB.

## Data and materials availability

All data are available in the main text or the supplementary materials. RNA sequencing data are deposited to the Gene Expression Omnibus, accession number GSE218686.

---

**Extended Data Fig. 1.**
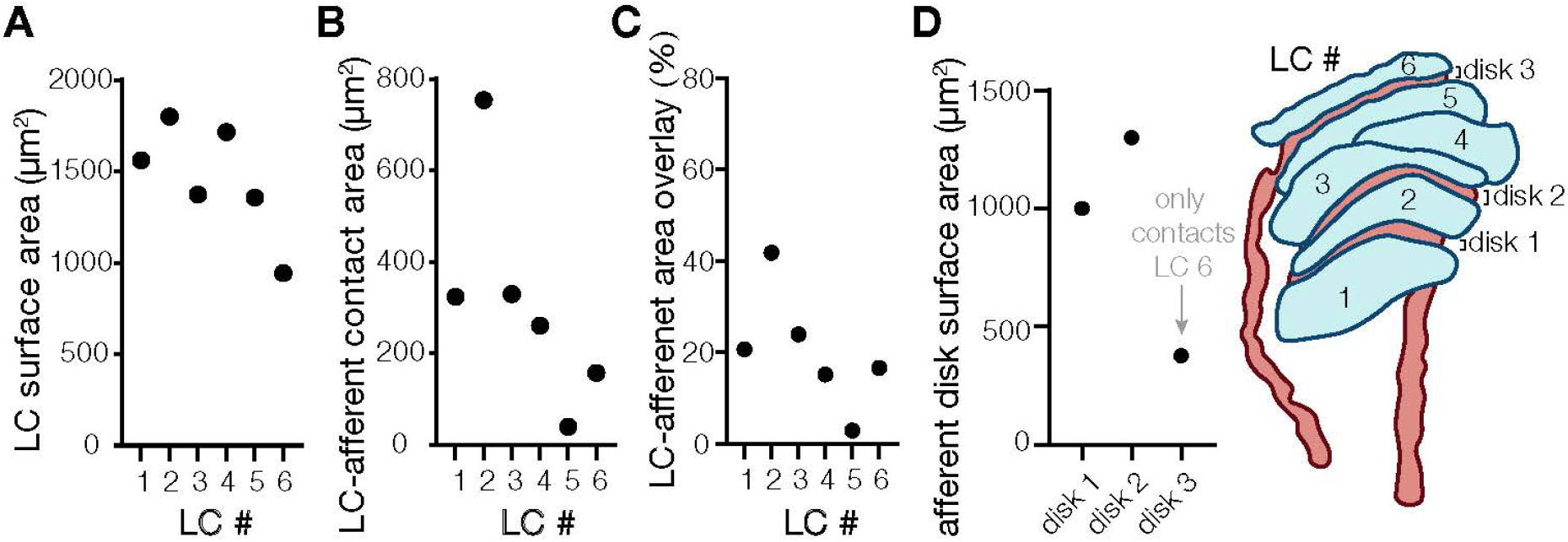
Quantification of lamellar cell – afferent contact area. **(A)** Quantification of the surface area of lamellar cells in a reconstituted avian Meissner corpuscle. **(B**) Quantification of LC surface area in contact with the afferent. **(C)** Quantification of the portion of LC surface area in contact with afferents. **(D)** Quantification of afferent disk area in contact with LCs. The cartoon depicts a dually innervated corpsucle.

**Extended Data Fig. 2.**
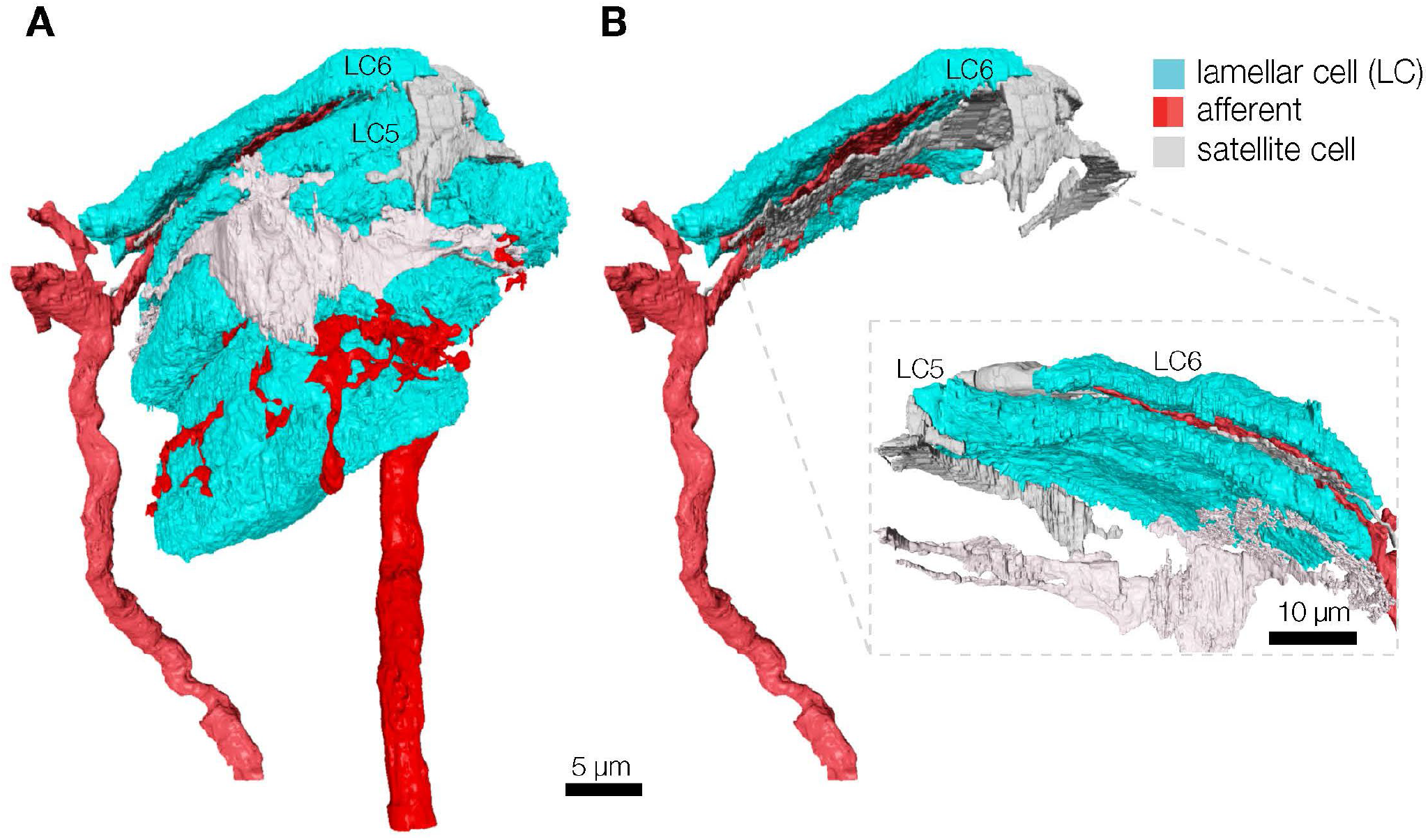
Three-dimensional image of a satellite cell in avian Meissner corpuscle. **(A)** 3D reconstitution of the corpuscle core with a satellite cell. **(B)** Isolated 3D images of satellite cell projections relative to LCs and the afferent.

**Extended Data Fig. 3.**
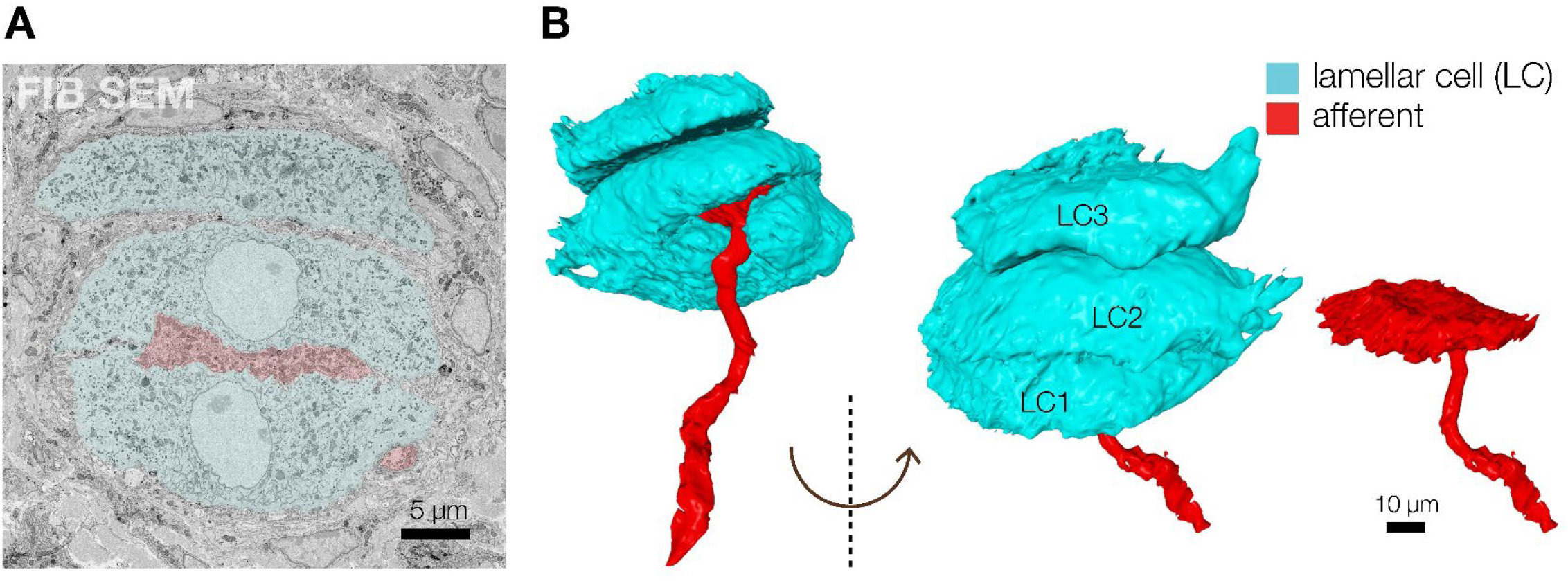
Three-dimensional image of an avian Meissner corpuscle innervated by a single afferent. **(A)** A pseudo-colored FIB-SEM image of a section of a Meissner corpuscle in duck bill skin with three LCs (blue) innervated by a single afferent (red). **(B)** Partial 3D reconstruction of a Meissner corpuscle core (without satellite cells) innervated by a single afferent. The afferent (red) forms a single disc positioned between LC1 and LC2 (blue).

**Extended Data Fig. 4.**
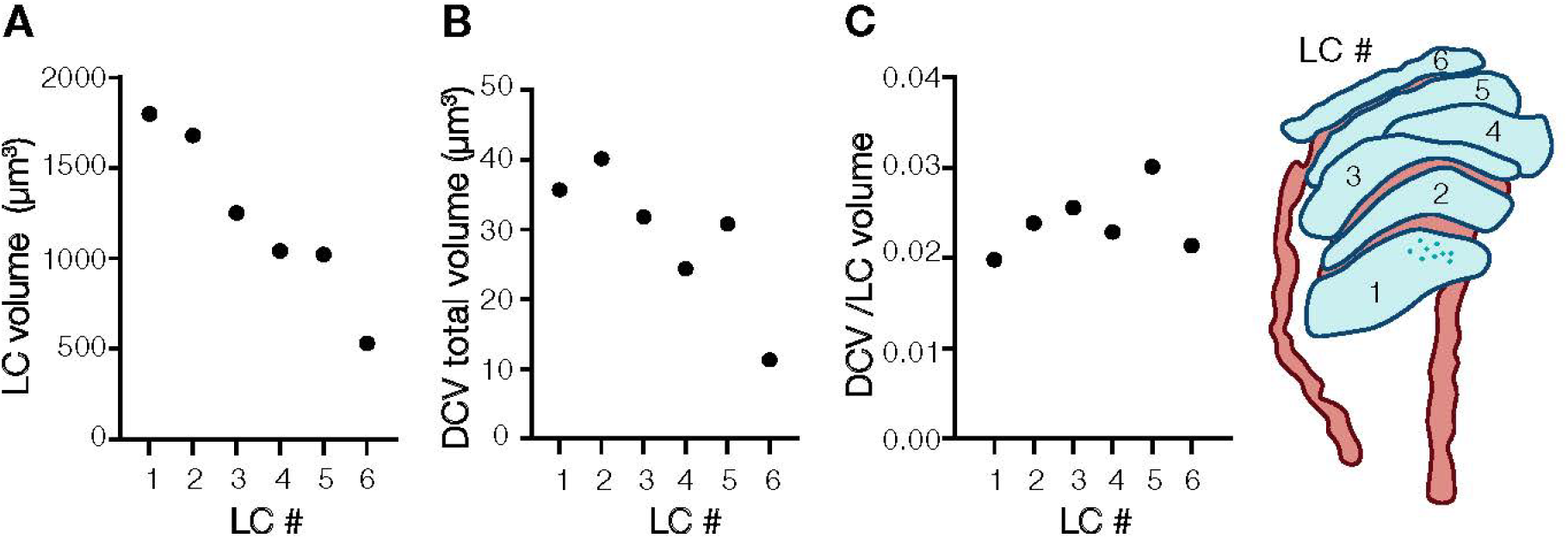
Quantification of dense core vesicle size and volume. **(A)** Quantification of the lamellar cell volume. **(B)** Quantification of the volume occupied by DCVs in lamellar cells. **(C)** Quantification of total DCV volume per LC. The cartoon depicts a dually innervated corpsucle.

**Extended Data Fig. 5.**
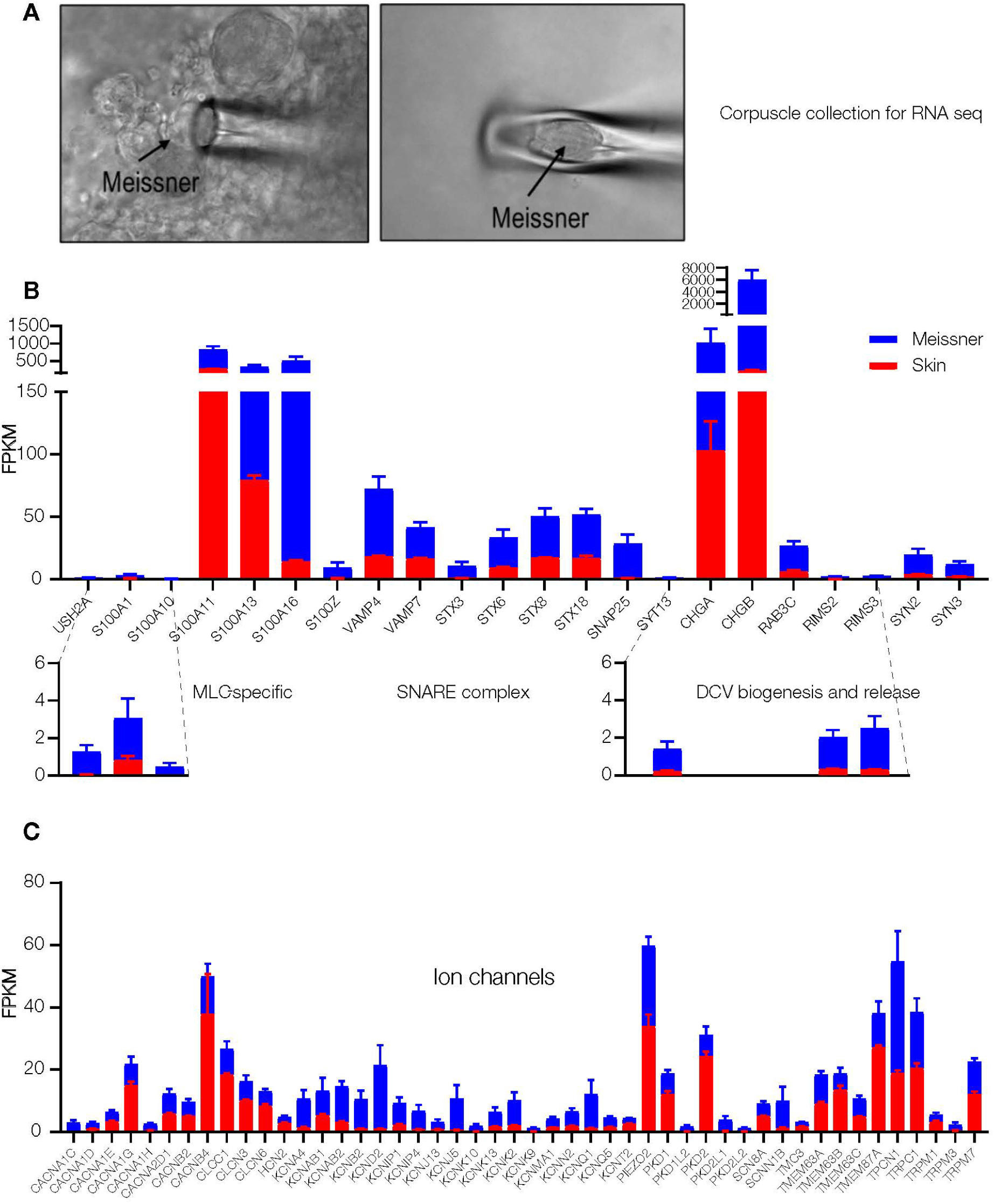
RNA-sequencing of avian Meissner corpuscles. **(A)** Images of Meissner corpuscles extracted from duck bill skin for RNA sequencing. **(B, C)** Quantification of transcript levels in Meissner corpuscles and adjacent skin areas. Data are mean ± SEM from 6 corpuscles and 5 skin samples. FPKM, fragments per kilobase of exon per million mapped fragments.

**Extended Data Fig. 6.**
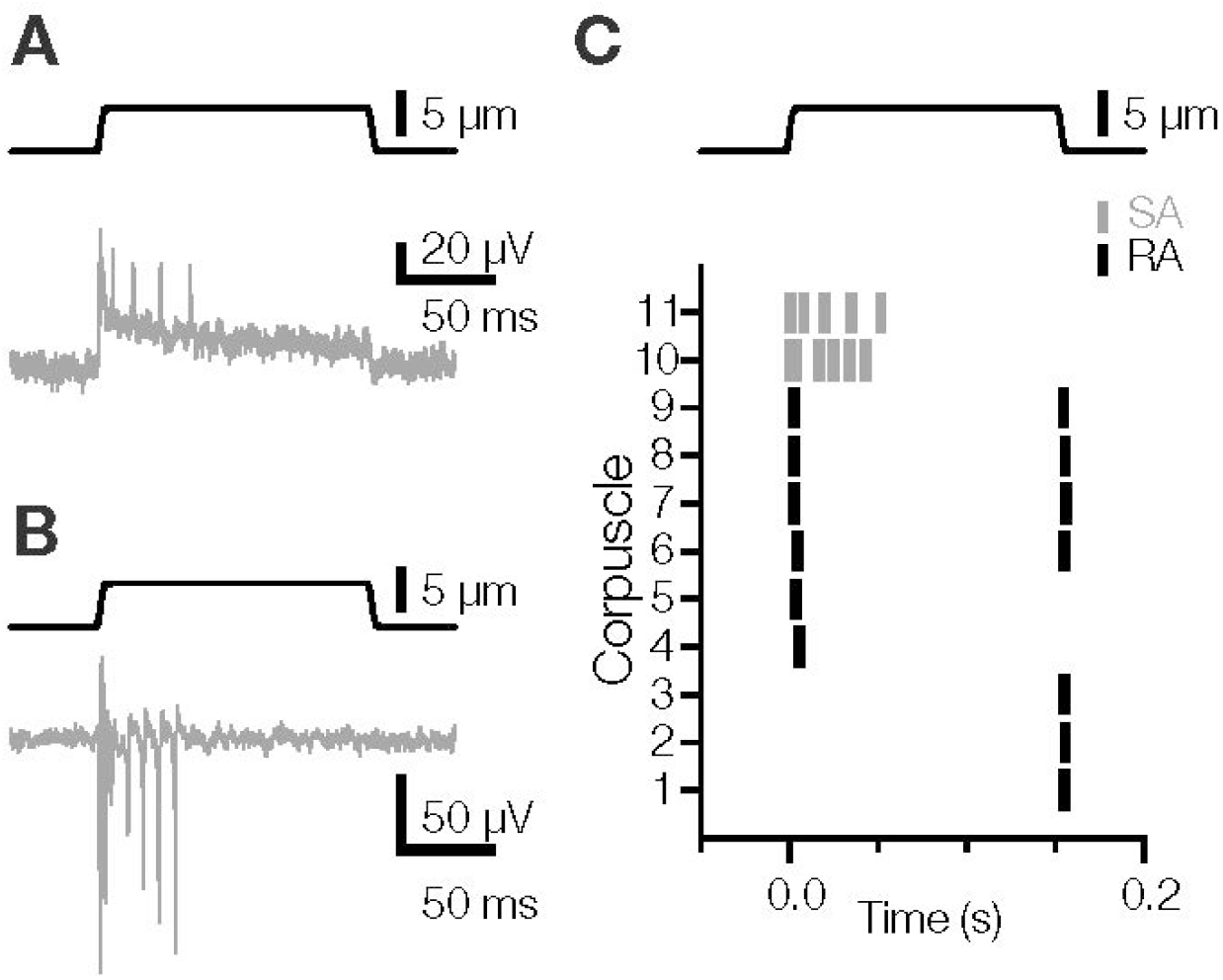
Slowly and rapidly adapting responses in the afferents from avian Meissner corpuscles. **(A)** Recordings of action potentials in the afferent in response to mechanical stimulation. **(B, C)** Raster plot of two slowly adapting afferents (SA) and representative rapidly afferents (RA) from different corpuscles in response to mechanical stimulation.

**Extended Data Fig. 7.**
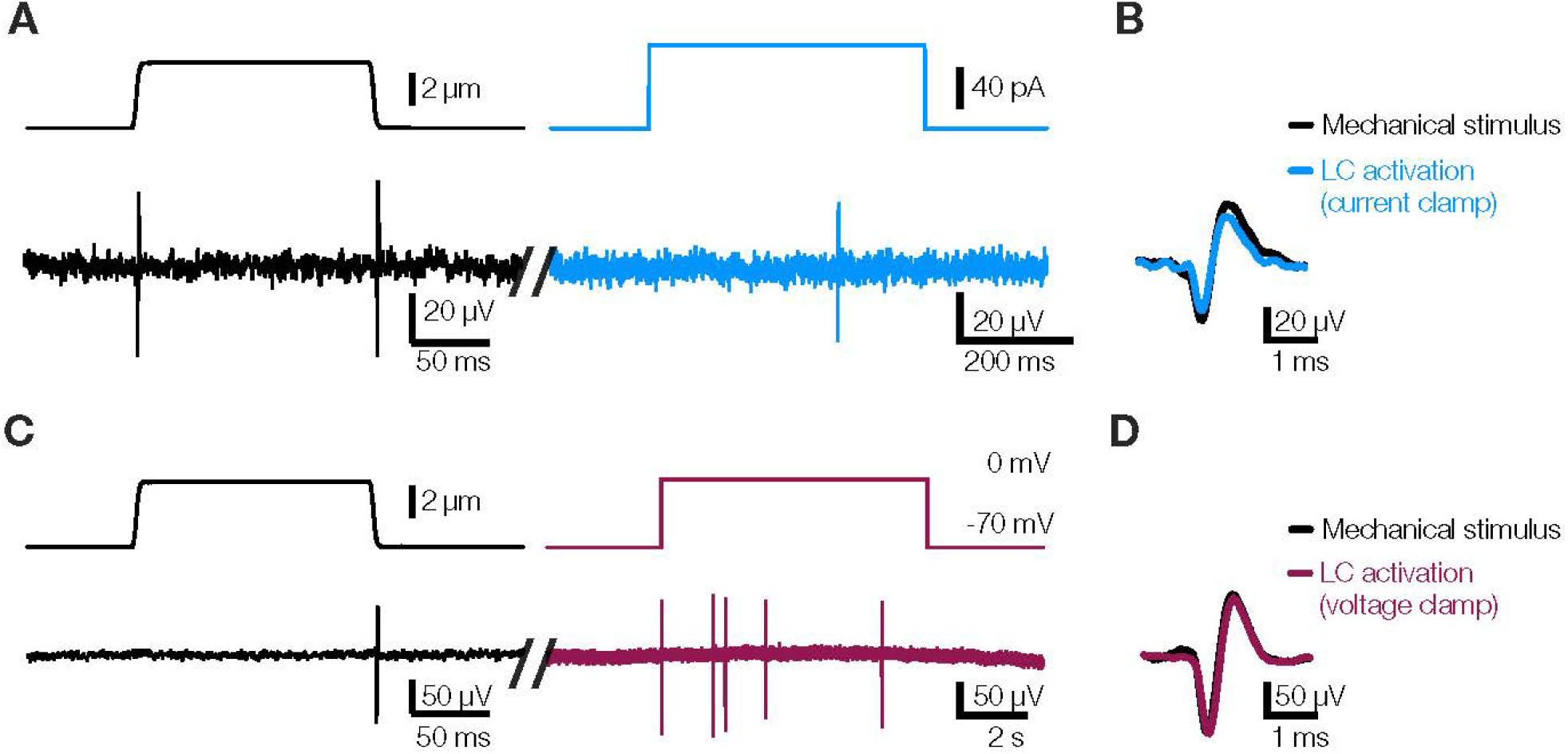
Action potentials evoked in the afferent by mechanical stimulation and LC activation are indistinguishable. (A) Exemplar traces recorded in a Meissner afferent in response to mechanical stimulation of the corpuscle (left) or LC activation by current injection. (B) Overlay of APs evoked by mechanical stimulation or LC activation by current injection. (C) Exemplar traces recorded in a Meissner afferent in response to mechanical stimulation of the corpuscle (left) or LC activation by depolarization. (D) Overlay of APs evoked by mechanical stimulation or LC activation by depolarization.

## Supplementary Information

**Supplementary Table I.** Accuracy of FIB-SEM data segmentation for a Meissner corpuscle innervated by two afferents.

| Object | IoU | F1 |
| --- | --- | --- |
| Afferents | 0.67 | 0.80 |
| DCVs | 0.80 | 0.89 |
| Lamellar cells | 0.99 | 0.99 |
| Satellite cells | 0.97 | 0.98 |
| Collagen | 0.94 | 0.97 |

**Video 1.** 3D architecture of a duck Meissner corpuscle innervated by two afferents (FIB-SEM).

**Video 2.** An image stack of a fragment of LC-afferent contact area (electron tomography). The video shows a cross-section of the afferent disk sandwiched between two DCV-containing LCs.

**Video 3.** 3D reconstruction of a fragment of LC-afferent contact area (electron tomography). Depicted are mitochondria (yellow), DCVs (red), LC membrane (dark blue), afferent membrane (purple), clear vesicles (light blue), caveolae (green), membrane densities resembling adherens junctions (orange).

**Video 4.** A close-up 3D reconstruction of a fragment of LC-afferent contact area (electron tomography), depicting fusing DCVs, a caveolae, and tethers connecting LC and afferent membranes.

## Materials and Methods

### Animals

Experiments with Mallard duck embryos (*Anas platyrhynchos domesticus*) were approved by and performed in accordance with guidelines of the Institutional Animal Case and Use Committee of Yale University (protocol 11526). Animals used in experiments were at development stages embryonic day 25 (E25) to E27, unless otherwise indicated.

### *Ex vivo* bill-skin preparation

Dissection of bill-skin was performed as described previously^13^. Briefly, the glabrous skin of the bill was quickly cut from the embryo in ice-cold L-15 media. The bill-skin was placed with the dermis on top and epidermis on the bottom in a recording chamber in Krebs solution containing (in mM) 117 NaCl, 3.5 KCl, 2.5 CaCl2, 1.2 MgCl2, 1.2 NaH2PO4, 25 NaHCO3, and 11 glucose, saturated with 95% O2 and 5% CO2 (pH 7.3-7.4), at room temperature (22-23°C). Corpuscles in the dermis were visualized on an Olympus BX51WI upright microscope with an ORCA-Flash 4.0 LT camera (Hamamatsu). The bill-skin was treated with 2 mg/mL collagenase P (Roche) in Krebs solution for 5 minutes, then washed with fresh Krebs solution before recording.

### Electrophysiology

#### Single-fiber recording

Recordings from single afferent fibers of avian Meissner (Grandry) corpuscles were acquired at room temperature in Krebs solution using a MultiClamp 700B amplifier and Digidata 1550A digitizer (Molecular Devices). Single-fiber recording pipettes were created from thin-wall, 1.5 mm diameter borosilicate glass capillaries. Pipettes were pulled using a P-1000 micropipette puller (Sutter Instruments) to create tip diameters of 5 to 30 μm, then filled with Krebs solution. Pipettes were placed on a CV-7B headstage connected to a High-Speed Pressure Clamp (ALA Scientific Instruments). Single corpuscles and connected afferents within the same field of view were identified under a 40X objective lens. The recording pipette was placed next to the afferent, and negative pressure was applied until a large section (∼5 μm) of the afferent was sucked into the pipette. The extracellular afferent voltage was recording in current-clamp mode, sampled at 20 kHz and low-pass filtered at 1 kHz. A suprathreshold mechanical step or 20 Hz vibrating stimulus was applied to the connected corpuscle to confirm the presence of mechanically induced action potentials in the afferent fiber. Fresh Krebs solution was regularly perfused onto the preparation between recordings.

Mechanical stimuli were applied to a single corpuscle using a blunt glass probe (5 to 10 μm tip diameter) mounted on a piezoelectric-driven actuator (Physik Instrumente Gmbh). A mechanical step stimulus was applied to corpuscles with variable displacements in increments of 1 μm. The duration of the static phase of the step stimulus was constant at 150 ms. The duration of the dynamic phases of the step stimulus was 3 ms or 30 ms to record single or multiple action potentials with the displacement ramp, respectively. Vibratory stimuli were applied using a 20 Hz sinusoidal waveform with a constant amplitude between 1 and 38 μm.

#### Patch-clamp electrophysiology

Recordings of lamellar cells were performed as described previously^3^ with simultaneous single-fiber recording in Krebs solution described above. Standard-wall, 1.5 mm diameter borosilicate pipettes with tip resistances of 2-4 MΩ were used to acquire lamellar cell voltage-clamp and current-clamp recordings. Pipettes were filled with (in mM) 135 K-gluconate, 5 KCl, 0.5 CaCl2, 2 MgCl2, 5 EGTA, 5 HEPES, 5 Na2ATP, and 0.5 Na2GTP (pH 7.3 with KOH) and placed on a secondary CV-7B headstage (connected to a second High Speed Pressure Clamp) which was placed perpendicular to the primary headstage. After single-fiber recordings were established for an individual corpuscle, the whole-cell recording configuration was achieved for one of the lamellar cells within the corpuscle. Lamellar cell current and voltage data was sampled at 20 kHz and low pass filtered at 2 kHz.

Lamellar cells were activated by current injection in current-clamp or a voltage step in voltage-clamp, while the afferent activity was measured simultaneously via single-fiber recording. In current-clamp, depolarizing current steps (from 10 to 120 pA in 10 pA increments) were applied to elicit voltage gated calcium channel-mediated action potentials in the lamellar cell. In voltage-clamp, the lamellar cell was initially held at -70 mV, then clamped at 0 mV for 10 seconds. Mechanical step stimuli were applied before and after lamellar cell activation protocols to elicit mechanoreceptor APs, confirming health and proper function of the corpuscle and afferent throughout the experiment. To test the potential mechanism of LC-afferent communication, lamellar cells were held at -70 mV, then voltage-clamped at 0 mV for 5 seconds while measuring afferent activity in each experimental condition. Intracellular solution containing (in mM) 133 CsCl, 10 HEPES, 5 EGTA, 1 CaCl_2_, 1 MgCl_2_, 4 MgATP, and 0.4 Na_2_GTP (pH = 7.3 with CsOH) was used in lamellar cells to block potassium current. To block calcium influx, Krebs solution containing 20 μM CaCl_2_ and 300 μM CdCl_2_ was added, then washed off with normal Krebs solution.

Single-fiber and patch-clamp recordings were acquired from individual corpuscles in skin preparations from different animals. Electrophysiological data was obtained in Clampfit 10.7 (Molecular Devices), then analyzed and displayed in GraphPad Prism 9.5.1 (GraphPad Software, LLC).

### FIB-SEM

#### EM sample preparation

The immersion fixation protocol was adapted from the mouse skin preparation method ^2^. A patch of bill skin was dissected from a duck embryo and immediately immersed in fixative solution containing 2.5% glutaraldehyde, 2.5% paraformaldehyde, 0.13M cacodylate, 4 mM CaCl2, 4 mM MgCl2 (pH 7.4, 37°C). The epidermis was removed while the skin patch remained in this solution, and the skin was cut into 1 mm by 1 mm sections at room temperature. After 2 hours, the dermis was transferred to fresh solution and gently shaken at 4°C for 48 hours. The solution was replaced with freshly prepared fixative at the 24 hour timepoint. After 48 hours, the sample was stored in a solution of 1.5% paraformaldehyde, 0.13M cacodylate pH 7.4 and stored at 4°C.

The bill skin samples were then sectioned into 300 µm thick slices in 0.13 M cacodylate buffer using a Compresstome (Precisionary, MA). The slices were washed in cacodylate buffer (0.13 M), postfixed with 2% osmium tetroxide and 1.5% potassium ferrocyanide in 0.13 M cacodylate buffer for 120 min at 0°C. After wash in distilled water, the slices were stained with 1% thiocarbohydrazide for 40 min at 40°C, 2% osmium tetroxide for 90 min at room temperature followed by 1% uranyl acetate at 4°C overnight. These staining reagents were diluted in the double distilled water. The sample slices were completely washed with distilled water between each step at room temperature three times for 10 min each. Finally, the slices were transferred into lead aspartate solution at 50°C for 120 min followed by distilled water wash at room temperature three times for 10 min each. After the heavy metal staining procedure, the samples were dehydrated with graded ethanol, embedded in Durcupan resin (Sigma, MO) and then polymerization at 60°C for 48 hours.

#### FIB-SEM sample preparation

One Durcupan embedded duck bill skin sample contained one whole Meissner corpuscle and one partial Meissner corpuscle (DB.B2-01M), and another sample contained one whole Meissner and one whole Pacinian corpuscle (DB.B2-01MP). The sample was first mounted to the top of a 1 mm copper post which was in contact with the metal-stained sample for better charge dissipation, as previously described^31^. The vertical sample post was trimmed to a small block containing one Meissner corpuscle with the width perpendicular to the ion beam, and the depth in the direction of the ion beam sequentially^32^. The block size was 110 x 80 µm^2^. After the entire volume of DB.B2-01M being FIB-SEM imaged, the sample post was then to trimmed to sample DB.B2-01MP that contains Region of Interest (ROI) of one Meissner conjunction with one Pacinian corpuscle (DB.B2-01MP) with block size of 135 x 110 µm^2^. The trimming was guided by X-ray tomography data obtained by a Zeiss Versa XRM-510 and optical inspection under a microtome. Thin layers of conductive material of 10-nm gold followed by 100-nm carbon were coated on the trimmed samples using a Gatan 682 High-Resolution Ion Beam Coater. The coating parameters were 6 keV, 200 nA on both argon gas plasma sources, 10 rpm sample rotation with 45-degree tilt.

#### FIB-SEM 3D large volume imaging

Two FIB-SEM prepared samples, DB.B2-01M and DB.B2-01MP were imaged sequentially by two customized Zeiss FIB-SEM systems previously described^31, 33, 34^. The block face of ROI was imaged by a 2 nA electron beam with 1.2 keV landing energy at 2 MHz scanning rate. The x-y pixel resolution was set at 8 nm. A subsequently applied focused Ga^+^ beam of 15 nA at 30 keV strafed across the top surface and ablated away 8 nm of the surface. The newly exposed surface was then imaged again. The ablation – imaging cycle continued about once every half minute for four days to complete FIB-SEM imaging DB.B2-01M that contains one Meissner corpuscle, and about once every minute for one week to complete DB.B2-01MP that contains one Meissner and one Pacinian corpuscles. The acquired image stack formed a raw imaged volume, followed by post processing of image registration and alignment using a Scale Invariant Feature Transform (SIFT) based algorithm. The aligned stack consists of a final isotropic volume of 60 x 50 x 58 µm^3^ and 85 x 56 x 75 µm^3^ for DB.B2-01M and DB.B2-01MP, respectively. The voxel size of 8 x 8 x 8 nm^3^ was maintained for both samples throughout entire volumes, which can be viewed in any arbitrary orientations.

#### FIB-SEM segmentation

The segmentation of organelles, cells, and nuclei from FIB-SEM images was achieved with Apeer, an AI-driven cloud-based platform (https://www.apeer.com/)^35^. Deep learning techniques were utilized to achieve automated segmentation, employing a customized convolutional neural network (CNN) architecture based on 3D U-Net. To generate ground truth data, cells and organelles were manually annotated from a small set (100 planes) of the raw FIB-SEM images. The CNNs were trained using the annotated ground truth data and proofread to achieve high-quality segmentation of the objects in 3D. Semantic segmentation was applied to each object, and the accuracy of the segmentation was assessed by evaluating the voxel Intersection over Union (IoU) and F1 scores. Apeer machine learning models were downloaded separately for each class of cells or organelles to create a full 3D model on a full dataset. All volumes were segmented at 16 x 16 x 16 nm resolution except DCVs which were segmented at 8 x 8 x 8 nm.

#### FIB-SEM reconstruction and data analysis

Raw FIB-SEM data and Apeer machine learning models for each class were imported to Arivis Vision 4D software. Using this software, each individual cell and organelle was segmented to generate complete objects, which were then filtered by size to remove any extraneous noise components. The objects were manually proofread and adjusted as necessary, and various quantitative measures such as volume, distances, surface area, and diameters were calculated within the software. The diameter of each DCV was estimated by identifying the longest shortest path between two mesh nodes. The contact area between the LC and the afferent was determined by dilating each object by 1 pixel, creating their intersection, and dividing the resulting surface area by two. Python scripts were utilized to calculate density scores for the DCVs, and videos were generated using Arivis Vision 4D. 3D tomography was reconstructed at 1.6 x 1.6 x 1.6 nm resolution manually per plane using Arivis Vision 4D software.

### 3D transmission electron microscopy tomography

Freshly peeled duck bill skin was fixed in Karnovsky fixative at 4°C for 1 hour, washed in 0.1 M sodium cacodylate buffer (pH 7.4), then postfixed in 1% osmium tetroxide for 1 hour in the dark on ice. The tissue was stained in Kellenberger solution for 1 hour at room temperature after washing in distilled water, dehydrated in a series of alcohols and propylene oxide, then embedded in EMbed 812, and polymerized overnight at 60°C. Thick sections of 250 nm depth were obtained from hardened blocks using a Leica UltraCut UC7 on copper formvar coated slot grids. 250 nm thick sections were contrast stained using 2% uranyl acetate and lead citrate and 15nm fiducial gold was added to both sides to aid alignment for Tomography. Sections were viewed using a FEI Tecnai TF20 at 200 Kv and data was collected using SerialEM^36^ on a FEI Eagle 4Kx4K CCD camera using tilt angles of -60 to 60 degrees, then reconstructed in IMOD (University of Colorado, Boulder). All solutions were supplied by Electron Microscopy Sciences (Hatfield, PA).

### Corpuscle RNA sequencing

Corpuscles were manually collected and pooled for subsequent transcriptomic analysis in RNase-free conditions. Aspiration pipettes pulled from capillary glass tubing (G150F-3, Warner Instruments, Hamden, CT) using a micropipette puller (P-1000, Sutter, Novato, CA) had a tip diameter of ∼40–60 µM, and were filled with 3 µl of the RNA Lysis Buffer (Quick-RNA Microprep Kit, Zymo, Irvine, Ca). The pipette was mounted on a micromanipulator and used to aspirate 2– 10 corpuscles from duck bill skin by applying negative pressure using a high-speed pressure clamp system (HSPC-1, ALA Scientific Instruments). Collected corpuscles were deposited into a 0.5 ml tube containing 10 µl of the RNA Lysis Buffer. Nearby skin cells devoid of any corpuscles were collected for comparison. Samples were then stored at −80°C until RNA isolation. 5-6 total replicates were collected from 3 independent skin isolations. RNA was isolated using the Quick-RNA Microprep Kit (Zymo) according to manufacturer’s instructions. RNA concentration and integrity number (RIN) were assessed by Agilent 2100 Bioanalyzer (Agilent, Santa Clara, CA). RNA concentrations for corpuscles were in the range of 63–323 pg/µl and RIN values were in the range of 6.1–8.0.

Library preparation and sequencing were carried out at the Yale Center for Genome Analysis. Sequencing libraries were prepared using the Kapa mRNA Hyper Prep kit (KAPA Biosystems, Wilmington, MA) (skin samples) or the NEBNext Single Cell/Low Input RNA Library Prep Kit (New England Biolabs, Ipswich, MA) (corpuscles samples). Libraries were sequenced on the Illumina NovaSeq instrument in the 150bp paired-end mode according to manufacturer’s protocols. A total of ∼30-52 million sequencing read pairs per sample were obtained.

The sequencing data was processed on the Yale Center for Research Computing cluster. Raw sequencing reads were filtered and trimmed to retain high-quality reads using Trimmomatic v0.39 ^37^ with default parameters. Filtered high-quality reads from all samples were aligned to the duck reference genome using the STAR aligner v2.7.7a with default parameters^38^. The duck reference genome and the gene annotation were obtained from the National Center for Biotechnology Information (Anas platyrhynchos; assembly ZJU1.0; NCBI Annotation Release: 104; all files accessed on 4/30/2021).

Reference genome:

https://ftp.ncbi.nlm.nih.gov/genomes/all/GCF/015/476/345/GCF_015476345.1_ZJU1.0/GCF_015476345.1_ZJU1.0_genomic.fna.gz;

Gene annotation:

https://ftp.ncbi.nlm.nih.gov/genomes/all/GCF/015/476/345/GCF_015476345.1_ZJU1.0/GCF_015476345.1_ZJU1.0_genomic.gff.gz;

The gene annotation was filtered to include only protein-coding genes. Aligned reads were counted by the featureCounts module within the Subread package v2.0.1 using the “unstranded” mode with default parameters ^39^. Read counting was performed at the gene level, i.e. the final read count for each gene included all reads mapped to all exons of the gene. Processing, normalization, and statistical analysis of read counts was performed using the EdgeR v3.38.4 package in R v4.2.1 ^40^. Normalized read counts were obtained by normalizing raw read counts to the effective library sizes of each sample. Effective library sizes were calculated by normalizing raw library sizes by RNA composition using a trimmed mean of M-values (TMM) method, as implemented in the calcNormFactors function of the EdgeR package. Normalized read counts were further normalized to gene length and expressed as “fragments per kilobase gene length per million mapped reads” (FPKM). Statistical analysis of differential expression of genes between groups was performed using the GLM approach and the quasi-likelihood F-test, as implemented in the EdgeR package. RNA sequencing data were deposited to the Gene Expression Omnibus, accession number GSE218686.

## References

1 Handler, A. & Ginty, D. D. The mechanosensory neurons of touch and their mechanisms of activation. Nat Rev Neurosci (2021). https://doi.org:10.1038/s41583-021-00489-x

2 Neubarth, N. L. et al. Meissner corpuscles and their spatially intermingled afferents underlie gentle touch perception. Science 368 (2020). https://doi.org:10.1126/science.abb2751

3 Nikolaev, Y. A. et al. Lamellar cells in Pacinian and Meissner corpuscles are touch sensors. Science advances 6 (2020). https://doi.org:10.1126/sciadv.abe6393

4 Schwaller, F. et al. USH2A is a Meissner’s corpuscle protein necessary for normal vibration sensing in mice and humans. Nature neuroscience 24, 74–81 (2021). https://doi.org:10.1038/s41593-020-00751-y

5 Ziolkowski, L. H., Gracheva, E. O. & Bagriantsev, S. N. Tactile sensation in birds: Physiological insights from avian mechanoreceptors. Current opinion in neurobiology 74, 102548 (2022). https://doi.org:10.1016/j.conb.2022.102548

6 Orefice, L. L. et al. Peripheral Mechanosensory Neuron Dysfunction Underlies Tactile and Behavioral Deficits in Mouse Models of ASDs. Cell 166, 299–313 (2016). https://doi.org:10.1016/j.cell.2016.05.033

7 Schneider, E. R., Gracheva, E. O. & Bagriantsev, S. N. Evolutionary Specialization of Tactile Perception in Vertebrates. Physiology 31, 193–200 (2016). https://doi.org:10.1152/physiol.00036.2015

8 Ranade, S. S. et al. Piezo2 is the major transducer of mechanical forces for touch sensation in mice. Nature 516, 121–125 (2014). https://doi.org:10.1038/nature13980

9 Kefauver, J. M., Ward, A. B. & Patapoutian, A. Discoveries in structure and physiology of mechanically activated ion channels. Nature 587, 567–576 (2020). https://doi.org:10.1038/s41586-020-2933-1

10 Abdo, H. et al. Specialized cutaneous Schwann cells initiate pain sensation. Science 365, 695–699 (2019). https://doi.org:10.1126/science.aax6452

11 Ojeda-Alonso, J. et al. Sensory Schwann cells are required for mechanical nociception and touch perception. bioRxiv, 2022.2002.2004.477749 (2022). https://doi.org:10.1101/2022.02.04.477749

12 Cobo, R. et al. in Somatosensory and Motor Research 1–15 (IntechOpen, 2020).

13 Ziolkowski, L. H., Gracheva, E. O. & Bagriantsev, S. N. Mechanotransduction events at the physiological site of touch detection. eLife 12, e84179 (2023). https://doi.org:10.7554/eLife.84179

14 Gottschaldt, K. M. The physiological basis of tactile sensibility in the beak of geese. J Comp Physiol 95, 29–47 (1974).

15 Gottschaldt, K. M. & Lausmann, S. Mechanoreceptors and their properties in the beak skin of geese (Anser anser). Brain research 65, 510–515 (1974).

16 Chang, W. et al. Merkel disc is a serotonergic synapse in the epidermis for transmitting tactile signals in mammals. Proc Natl Acad Sci U S A 113, E5491–5500 (2016). https://doi.org:10.1073/pnas.1610176113

17 Hoffman, B. U. et al. Merkel Cells Activate Sensory Neural Pathways through Adrenergic Synapses. Neuron 100, 1401–1413 e1406 (2018). https://doi.org:10.1016/j.neuron.2018.10.034

18 Hu, J., Chiang, L. Y., Koch, M. & Lewin, G. R. Evidence for a protein tether involved in somatic touch. EMBO J 29, 855–867 (2010). https://doi.org:10.1038/emboj.2009.398

19 Li, L. & Ginty, D. D. The structure and organization of lanceolate mechanosensory complexes at mouse hair follicles. eLife 3, e01901 (2014). https://doi.org:10.7554/eLife.01901

20 Garcia-Mesa, Y. et al. The acquisition of mechanoreceptive competence by human digital Merkel cells and sensory corpuscles during development: An immunohistochemical study of PIEZO2. Annals of anatomy = Anatomischer Anzeiger : official organ of the Anatomische Gesellschaft 243, 151953 (2022). https://doi.org:10.1016/j.aanat.2022.151953

21 Garcia-Mesa, Y. et al. Merkel cells and Meissner’s corpuscles in human digital skin display Piezo2 immunoreactivity. Journal of anatomy (2017). https://doi.org:10.1111/joa.12688

22 Berkhoudt, H. The morphology and distribution of cutaneous mechanoreceptors (Herbst and Grandry corpuscles) in bill and tongue of the Mallard (Anas platyrhynchos L.). Neth J Zool 30, 1–34 (1980).

23 Schneider, E. R. et al. Molecular basis of tactile specialization in the duck bill. Proc Natl Acad Sci U S A 114, 13036–13041 (2017). https://doi.org:10.1073/pnas.1708793114

24 West, A. K. et al. Quantitative Evaluation of Tactile Foraging Behavior in Pekin and Muscovy Ducks. Front Physiol 13, 921657 (2022). https://doi.org:10.3389/fphys.2022.921657

25. Zweers, G. A. Mechanics of the Feeding of the Mallard (Anas Platyrhynchos, L.; Aves, Anseriformes). 1St Edition edn, (S Karger Pub, 1977).

26 Handler, A. et al. Three-dimensional reconstructions of mechanosensory end organs suggest a unifying mechanism underlying dynamic, light touch. bioRxiv, 2023.2003.2017.533188 (2023). https://doi.org:10.1101/2023.03.17.533188

27 Ide, C. & Munger, B. L. A cytologic study of Grandry corpuscle development in chicken toe skin. J Comp Neurol 179, 301–324 (1978). https://doi.org:10.1002/cne.901790205

28 Ikeda, R. et al. Merkel cells transduce and encode tactile stimuli to drive abeta-afferent impulses. Cell 157, 664–675 (2014). https://doi.org:10.1016/j.cell.2014.02.026

29 Maksimovic, S. et al. Epidermal Merkel cells are mechanosensory cells that tune mammalian touch receptors. Nature 509, 617–621 (2014). https://doi.org:10.1038/nature13250

30 Woo, S. H. et al. Piezo2 is required for Merkel-cell mechanotransduction. Nature 509, 622–626 (2014). https://doi.org:10.1038/nature13251

31 Xu, C. S. et al. Enhanced FIB-SEM systems for large-volume 3D imaging. eLife 6 (2017). https://doi.org:10.7554/eLife.25916

32. Pang, S. & Xu, C. S. in Methods in Cell Biology: Volume Electron Microscopy Vol. 181 (ed P. Verkade) (Elsevier, (in press)).

33 Xu, C. S. et al. An open-access volume electron microscopy atlas of whole cells and tissues. Nature 599, 147–151 (2021). https://doi.org:10.1038/s41586-021-03992-4

34 Xu, C. S., Hayworth, K. J. & Hess, H. F. Enhanced FIB-SEM systems for large-volume 3D imaging. US Patent 10,600,615 B2 (2020).

35 Dang, D. et al. APEER: An Interactive Cloud Platform for Microscopists to Easily Deploy Deep Learning. Zenodo (2021). https://doi.org:10.5281/zenodo.5539895

36 Mastronarde, D. N. Automated electron microscope tomography using robust prediction of specimen movements. J Struct Biol 152, 36–51 (2005). https://doi.org:10.1016/j.jsb.2005.07.007

37 Bolger, A. M., Lohse, M. & Usadel, B. Trimmomatic: a flexible trimmer for Illumina sequence data. Bioinformatics 30, 2114–2120 (2014). https://doi.org:10.1093/bioinformatics/btu170

38 Dobin, A. et al. STAR: ultrafast universal RNA-seq aligner. Bioinformatics 29, 15–21 (2013). https://doi.org:10.1093/bioinformatics/bts635

39 Liao, Y., Smyth, G. K. & Shi, W. featureCounts: an efficient general purpose program for assigning sequence reads to genomic features. Bioinformatics 30, 923–930 (2014). https://doi.org:10.1093/bioinformatics/btt656

40 Robinson, M. D., McCarthy, D. J. & Smyth, G. K. edgeR: a Bioconductor package for differential expression analysis of digital gene expression data. Bioinformatics 26, 139–140 (2010). https://doi.org:10.1093/bioinformatics/btp616

